# SARS-CoV-2 3CLPro Dihedral Angles Reveal Allosteric Signaling

**DOI:** 10.1101/2024.05.22.595309

**Authors:** Daniel Evans, Samreen Sheraz, Albert Lau

## Abstract

In allosteric proteins, identifying the pathways that signals take from allosteric ligand-binding sites to enzyme active sites or binding pockets and interfaces remains challenging. This avenue of research is motivated by the goals of understanding particular macromolecular systems of interest and creating general methods for their study. An especially important protein that is the subject of many investigations in allostery is the SARS-CoV-2 main protease (Mpro), which is necessary for coronaviral replication. It is both an attractive drug target and, due to intense interest in it for the development of pharmaceutical compounds, a gauge of the state-of-the-art approaches in studying protein inhibition. Here we develop a computational method for characterizing protein allostery and use it to study Mpro. We propose a role of the protein’s C-terminal tail in allosteric modulation and warn of unintuitive traps that can plague studies of the role of protein dihedrals angles in transmitting allosteric signals.

## Introduction

Protein allostery is the phenomenon by which ligand binding at one site on a protein influences binding or enzymatic catalysis at another, possibly distant, site. Studies of allostery have evolved from the pioneering work of Monod-Wyman-Changeux^1^ and Koshland-Némethy-Filmer^2^ in defining the concerted MWC and sequential KMF models, respectively, of oxygen binding to hemoglobin. More recent work includes expanding the MWC model^3^, discussing the phenomenon of “dynamic allostery” without visible conformational change^4,5,6^, and studying allostery in intrinsically disordered proteins^7^. Techniques for understanding and controlling allostery have been developed through a combination of experiment and computation^8^. As these techniques become available, the power of allosteric drugs is becoming increasingly clear^9^. Fundamental questions about allostery, however, remain unanswered. The construction and testing of general models of allostery is an ongoing challenge.^10^ The abundance of allosteric pathways^11^ and druggable sites ^12,13^ and the accuracy of computational techniques for studying allostery^14,15^ remain the subject of debate. There is great demand for elucidating the molecular mechanisms governing allosteric processes.

A noteworthy example of an allosterically regulated protein is the SARS-CoV-2 main protease (Mpro). This protein is essential for viral replication, making it a well-studied, albeit incompletely understood, drug target. Mpro functions as a homodimer^16^, and there is evidence that the two subunits adopt different conformations^17,18,19^. The locations of ligand-binding sites on Mpro have been studied extensively^20,21,22,23,24,25^. Similarly, Mpro allosteric mechanisms have been explored by employing approaches such as crystal structure analysis^26^, elastic network modeling^27^, and molecular dynamics (MD) simulations^28^. Although Mpro inhibitors exist (including the FDA-approved nirmatrelvir), the continuing threat of COVID-19 infection motivates the search for better Mpro inhibitors.

Even for known allosteric Mpro inhibitors, determining the exact allosteric mechanism can be difficult. This is exemplified in the case of the inhibitor called x1187. Its binding site is known^23^, and it has been shown to inhibit dimerization^29^. However, a substantial percentage of Mpro is still a dimer even at high concentrations of x1187^29^. The biological assembly of the x1187-Mpro crystal is a symmetric homodimer, indicating that x1187 does not incur large-scale steric clashes that would prevent dimerization^23^. It has been suggested that x1187 might have additional mechanisms of inhibiting Mpro activity besides inhibiting dimerization^29^. The atom-level details of how x1187 inhibits Mpro, including disruption of dimerization and possible additional conformational changes, remain unclear.

One method for studying allostery is by examining how dihedral angles behave during MD simulations. If an allosteric site’s conformation influences an active site’s conformation, then there should be relatedness between dihedrals at the allosteric and active sites. This type of analysis is rather common. However, it has proven difficult to distinguish genuine dependencies between dihedrals from statistical noise and artifacts^14,15^. Additionally, the availability of many different statistical techniques, and their tendency to produce differing results^15,30^, makes it hard to interpret and compare these analyses. Clarity about when and how dihedral relatedness in MD simulations can be viewed as evidence of genuine allosteric communication is needed.

Here we propose a mechanism by which x1187 inhibits dimerized Mpro. In the process, we developed new software that can be used to study allostery in other proteins. We also found insights about how to interpret and judge studies of dihedral relatedness.

## Results

We studied Mpro allostery using previously released molecular dynamics (MD) simulation data^31^. To facilitate our analyses, we developed a new Python package called EVADE (Evaluation of Volume And Dihedral Exploration). EVADE contains two key capabilities. The first is the calculation of binding pocket volumes in an MD trajectory. Volume calculation uses a voxelized procedure based on POVME^32,33,34^. Our implementation includes a unique Jupyter visualization process (Figure 1), and uses MDAnalysis^35,36^ to increase flexibility of input formats. The second key capability of EVADE is the calculation of dihedral angles and relatedness between them. EVADE can compare dihedrals using measures including mutual information, correlation, covariance, and inverse covariance. EVADE is available at https://github.com/laulab-johnshopkins/evade.git.

**Figure 1.**
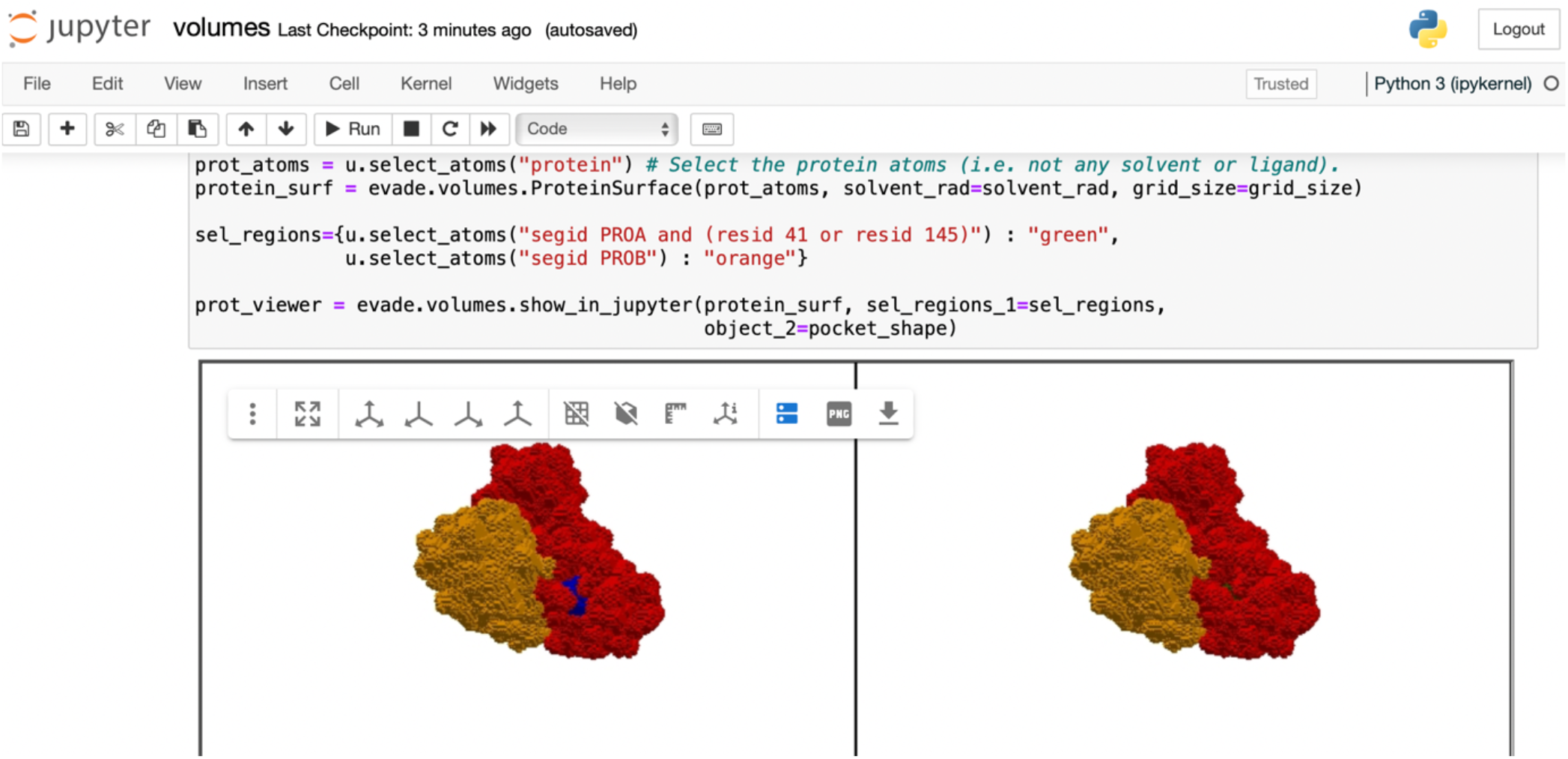
A screenshot of EVADE’s Jupyter notebook interface. The two subunits of the protein are shown in red and orange. The active site is shown in blue.

We began by looking for evidence of Mpro undergoing conformational change. The crystal structure of Mpro bound to the allosteric inhibitor x1187 is very similar to the structure of Mpro with a ligand in the active site (PDB IDs 5RFA^23^ and 6Y2F^18^). The most striking differences are that the allosterically inhibited structure is missing a model of the C-terminal tail (Figure 2A), presumably due to conformational dynamics, and undergoes sidechain rearrangement at the allosteric site (Figure 2B). Further evidence that Mpro is relatively rigid except for the C-terminal tail is provided by a 100-μs MD simulation from D.E. Shaw Research^31^. In agreement with data from other simulations^37^, the D.E. Shaw trajectory shows low root mean square fluctuations (RMSF) outside of the C-terminal tail (Figure 2C). Therefore, we inferred that we should focus on small-scale motions throughout the protein and larger motions at the C-terminal tail.

**Figure 2.**
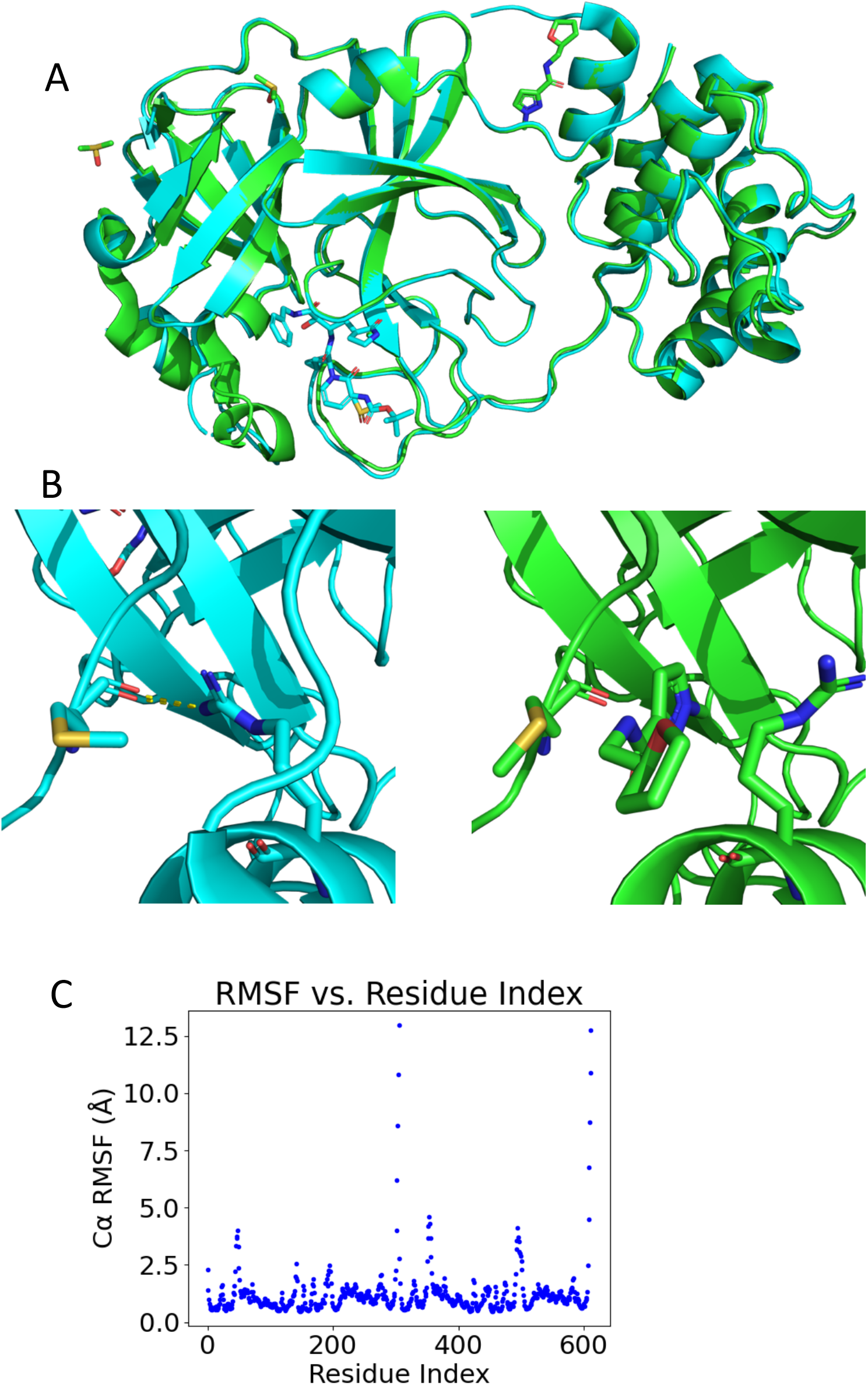
Most of Mpro is somewhat rigid. (A) Alignment of Mpro bound to a competitive inhibitor (6Y2F; blue) and an allosteric inhibitor (5RFA; green). The allosteric inhibitor is at the bottom. The C-terminal tail is at the top. The biological assembly is a homodimer. (B) The allosteric site in the absence of ligand (6Y2F; blue) and bound to x1187 (5RFA; green). Residues 6 and 298 are shown in sticks. Without x1187, residues 6 and 298 form a hydrogen bond (yellow line on left). X1187 pushes residue 298 away and breaks the hydrogen bond. (C) Alpha carbon RMSF vs. residue index. The two regions of high RMSF are the C-terminal tails of each protomer.

We used EVADE to investigate whether the C-terminal tail has the potential to be involved in allosteric regulation. The C-terminal tail fits into the active site and thus can be an inhibitor^38,39^ of enzyme function. Previous analysis of the D.E. Shaw trajectory reported that the tail spends some time at the active site^40^, but the implications of this conformation are not discussed. We were surprised to find that the tail binds the active site and remains there for roughly 14 *μ*s during the 100 *μ*s simulation, in a way that would present steric hindrance to substrate binding. The tail’s occupation of the active site is stabilized by a polar contact between the backbone of V303 and the sidechain of N119. Formation of this contact coincides with a major reduction in active site volume (Figure 3), from roughly 550 Å^3^ to roughly 350 Å^3^. The tail’s ability to block the active site even without ligand suggests that the tail might, when influenced by the right allosteric ligand, contribute to inhibition.

**Figure 3.**
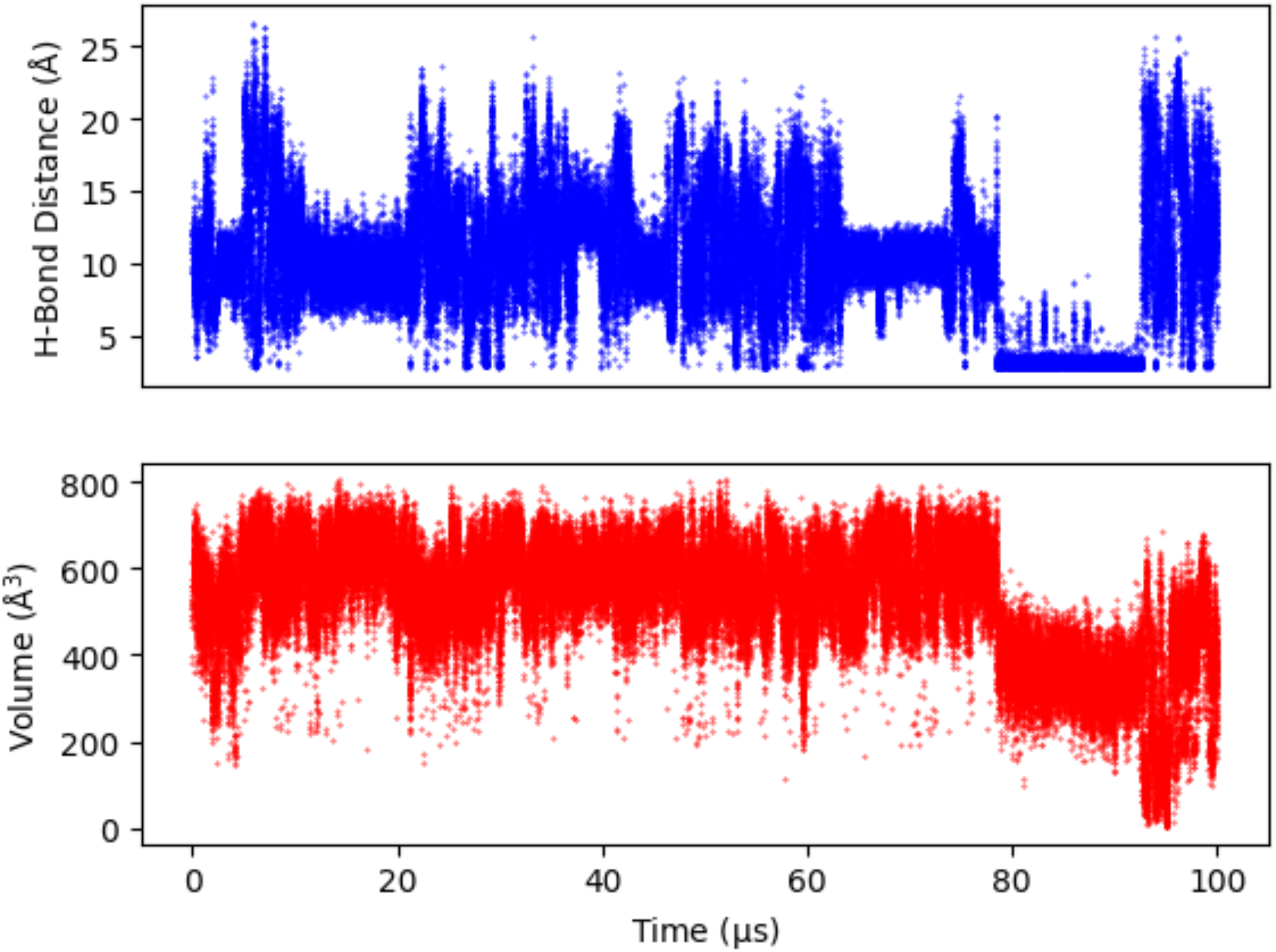
The C-terminal tail of chain A blocks the active site of chain B. Top: Hydrogen bond distance between chain A residue 303 and chain B residue 119. Bottom: volume of the chain B active site. Volume was calculated in EVADE, with the 6Y2F ligand used to de?ne the binding site.

Quantifying the propagation of allosteric signals through dihedrals requires distinguishing genuine allosteric communication from random dynamical fluctuations, i.e., noise. We used Mpro to test how susceptible to noise various measures of dihedral relatedness are. Definitions of statistical measures are in the supplement. For each pair of dihedrals in the protein, we studied how much the values of the two dihedrals are related to each other. We repeated this analysis for the first and second halves of the D.E. Shaw trajectory. Consistent with results for other proteins^15^, the inverse covariances determined from the first and second halves agree closely (Figure 4A). However, for many dihedral pairs, their correlation in the first half differs substantially from their correlation in the second half (Figure 4B). Such results were suggested to indicate that networks of dihedral correlations are irreproducible and should be viewed with caution^15^. However, we find that (at least for this system) differences in dihedral correlation between the first and second halves often reflect genuine differences in protein behavior between the halves. An example case is shown in Figure 4C, and a general trend is shown in Figure 4D. Disagreement between dihedral analyses on parallel MD datasets can indicate differences in the simulations themselves and are not necessarily indications of flawed analysis methodology.

**Figure 4.**
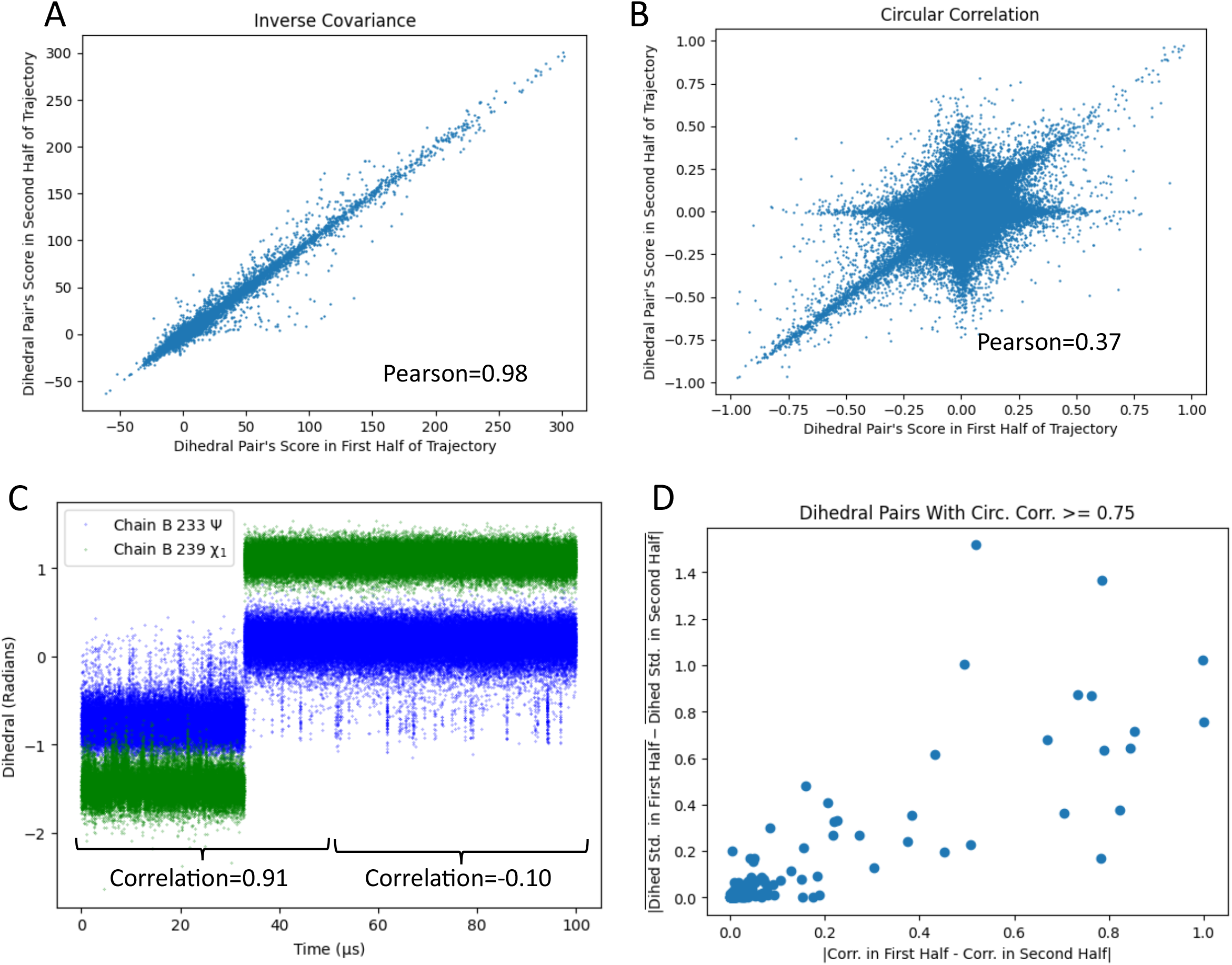
Dihedrals behave differently in the two halves of the DE Shaw trajectory. (A) Inverse covariances between pairs of dihedrals. Each point represents a pair of dihedrals. The x-axis is the inverse covariance between the dihedrals during the ?rst half of the trajectory. The y-axis is the inverse covariance between dihedrals during the second half of the trajectory. The graph resembles a line at y=x, indicating that results from the two halves are similar to each other. (B) The same as (A), but using correlation instead of inverse covariance. The ?rst half of the trajectory gives different results than the second half. (C) An example dihedral pair that gives different results for the ?rst and second half. The dihedrals appear strongly correlated when viewing the ?rst half. But when viewing only the second half, the correlation is not visible. (D) A graph showing that when correlations differ between trajectory halves, this is usually because the dihedrals themselves behave very differently.

How can we quantify the meaningfulness of dihedral relatedness scores calculated from MD simulations? While we cannot definitively answer this question, we can warn of factors that might bias such assessments. We suggest that observing similar scores in parallel datasets is a strong indication of veracity. However, comparing results from a subset of data against results from the entire dataset, which is sometimes done, requires caution. Including the same trajectory in both samples being compared can artificially increase calculated agreement (Table S1). Another source of caution involves the shape of correlation matrices. The elements where row and column indices are equal (i.e., diagonal elements) are a dihedral’s correlation with itself. In the case of Pearson correlation, the diagonals are all 1. For other metrics (e.g., mutual information), the diagonal values vary but tend to be high. In many cases, the diagonal elements are not useful (since correlation *between* residues is being studied). Including diagonals in reproducibility analyses can substantially inflate estimated reproducibility (Figure S1). While we are not aware of cases where diagonals were unfairly included when comparing two datasets, this mistake is easy enough to make that it deserves warning. Our observations do not negate the value of previously published dihedral analyses, which have been effective and rigorous. Rather, they provide guidance for ensuring that future studies continue this tradition of rigor.

We looked for evidence of mechanisms by which the x1187 binding site might allosterically inhibit Mpro. We noticed that the x1187 site is near the C-terminal tail, which (as discussed above) can block the active site. Using mutual information, we found that the x1187 site is allosterically connected, through a network of dihedral angle fluctuations, to the tail (Figure 5). To confirm that this is a genuine relationship, rather than being a result of statistical noise, we re-ran our analysis on another Mpro MD trajectory from Riken^41^ (Figure S2). In both the D.E. Shaw trajectory and the Riken trajectory, one protomer clearly shows allosteric connection between x1187 site and tail. Further work is needed to confirm the exact nature of the relationship between the x1187 allosteric site, the C-terminal tail, and Mpro activity.

**Figure 5.**
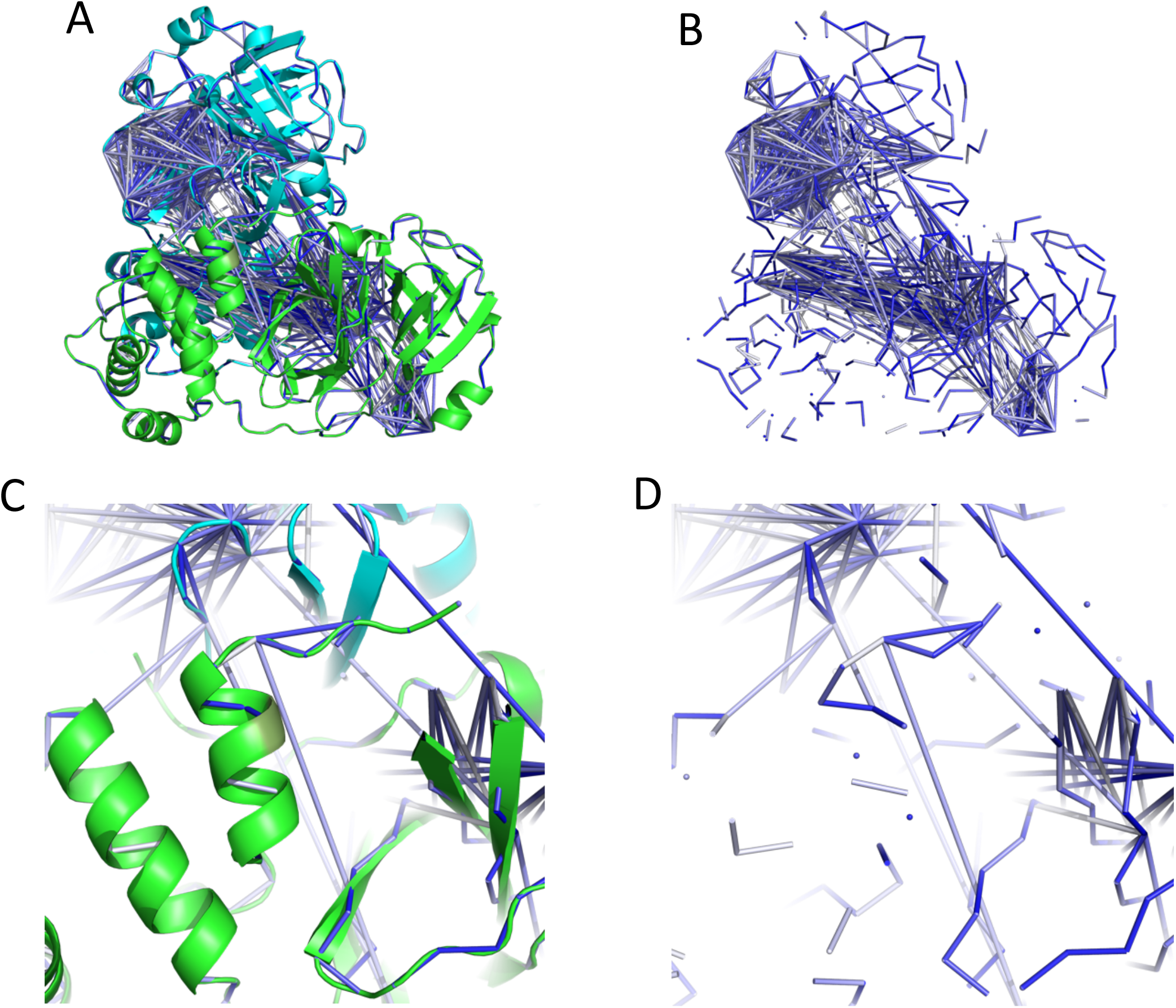
The x1187 site is allosterically coupled to the C-terminal tail. (A) An image of the Mpro dimer. The top 0.1% of dihedral pairs, as quanti?ed by mutual information, are shown as lines. Residue 298, which is involved in x1187 binding, is shown in dark green. (B) The dihedral pairs from (A), but without the protein. (C) A close-up of the C-terminal tail. As in (A), residue 298 is dark green. (D) Same as (C), but without the protein. This ?gure uses results from the D.E. Shaw simulation.

## Discussion

We studied dihedral-mediated allostery as both a generalizable phenomenon and a controller of Mpro function. We began by developing the EVADE software, which includes both a pocket volume calculator inspired by POVME^32,33,34^ and dihedral relatedness calculations. We used EVADE to evaluate whether dihedral correlations in MD simulations reveal genuine allostery or merely noise. We showed that when different datasets (in this case segments of a long trajectory) give different dihedral correlations, this does not necessarily indicate a problem with the relatedness calculation method. We also demonstrated flaws that should be avoided when assessing dihedral relatedness reproducibility. Lastly, we used our methods to study allostery in Mpro. We found that a previously reported allosteric site influences the conformation of the C-terminal tail, which can block the active site.

The allosteric site and the C-terminal tail, which we propose form a connected network, have both been previously (albeit incompletely) studied. The allosteric site has been crystallized with a ligand called x1187^23^, which has been shown to inhibit protein activity and dimerization^29^. However, the allosteric communication pathway is unknown, and there may be additional mechanisms of inhibition besides preventing dimerization^29^. The C-terminal tail has not been previously implicated in x1187 inhibition, but its interaction with the active site has been studied in the context of SARS-CoV-2 polyprotein cleavage^39^. Viral replication requires that Mpro have a C-terminal tail that is sterically compatible with its active site. Under normal conditions, Mpro’s C-terminal tail has too low an affinity for the active site to act as an autoinhibitor^38^; we suggest that x1187 could encourage active site-tail binding.

Our EVADE pocket volume calculator can be compared with other pocket-volume programs. Here we focus on programs where the user explicitly defines the region containing the pocket. The simplicity of this approach provides easily interpretable results. A notable example is POVME^32,33^. Users must provide PDB-format trajectories and define pockets by describing a set of inclusion/exclusion shapes. POVME3^34^ allows pockets to be defined by ligand shapes, but this version of POVME appears to be no longer maintained at the time of this writing. Another important example is EPOCK^42^, which (like EVADE) is inspired by POVME. EPOCK’s VMD plugin can take a variety of input formats, and it includes a user-friendly visualization process. EVADE includes features from both programs: it accepts many trajectory formats and has a built-in display of pocket regions (like EPOCK), and it can use ligand structures to define the pocket (like POVME3). EVADE also includes procedures for the necessary pre-processing step of aligning the input trajectory. Therefore, the EVADE volume finder alone is a uniquely useful tool.

The EVADE dihedral analysis code can also be compared with similar programs.

Programs for calculating dihedral relatedness include MDEntropy^43^, CARDS^44^, MDiGest ^30^, MutInf^45^, Allosteer^46^, and a Jupyter notebook for inverse covariance. MDEntropy and MutInf do not appear to be maintained as of this writing. MDiGest is recent and has an impressive set of features, although it does not yet include inverse covariance. CARDS is also an impressive software package. Although it only supports mutual information, it performs a relatively elaborate analysis that includes classification of when dihedral angles adopt a stable value vs. a disordered mix of values. EVADE presents a combination of features not found in any other dihedral analysis software.

We used EVADE to examine the consistency between datasets of calculated dihedral correlations. We found that dihedral correlations can be consistent across datasets, but careful analysis is needed to avoid biasing estimates of this. These statistical considerations are not merely of interest to developers of dihedral analysis methods. Rather, they are vital to anyone who evaluates studies of dihedral correlations. Our results are part of a broader conversation around the validity of dihedral correlation analysis^14,15^. We suggest that comparing results from different trajectories is an appropriate way of showing validity of dihedral correlations. Future work is needed to make assessment of agreement between trajectories more quantitative.

In conclusion, our paper presents results relevant to both the general study of dihedral correlations and the specific case of Mpro. We present new software, use it to assess consistency of dihedral correlations, and find a novel allosteric pathway in Mpro.

## Methods

EVADE is implemented in Python. Trajectory reading, alignment, and dihedral calculation are performed using MDAnalysis^35,36^ and its dependencies^47,48^. Pocket volumes are calculated using a voxelized approach described below. Voxel calculations are performed using trimesh^49^. Voxelized surfaces are displayed using PyVista^50^.

Pocket volumes are calculated using a similar procedure as in POVME^32,33,34^ and EPOCK^42^. The protein is first converted to a grid of cubes called voxels. Users provide a list of shapes that defines the binding site. The parts of the user-defined region that do not overlap with the protein are considered to be the binding pocket. This type of calculation requires that the trajectory be aligned. EVADE includes a function for aligning the trajectory based on only the pocket, which is preferable to aligning based on the whole protein^33^. EVADE also includes a Jupyter visualization process to help users check that their binding site is being defined correctly (Figure 1). Volumes calculated by EVADE are virtually identical to those from POVME (Figures S3a and S3b), demonstrating consistency in the two approaches.

Dihedral covariances and inverse covariances are calculated using established procedures^15^. Dihedral mutual information is calculated by placing values into 25 bins between –pi and pi. Results show general agreement with CARDS^44^ (Figure S3c), despite differences in calculation methodology.

Most of our analyses used a 100-μs MD simulation of the Mpro dimer provided by D.E. Shaw^31^. The exception is when we confirmed the allosteric connection between x1187 site and tail using a 10-μs trajectory provided by Riken^41^.

EVADE is available at https://github.com/laulab-johnshopkins/evade.git.

## Author Contributions

D.J.E. conceptualized the project, developed software, analyzed data, and wrote the manuscript. S.S. analyzed data and discovered concepts related to dihedral consistency and reviewed the manuscript. A.L. conceptualized the project, analyzed data, and wrote the manuscript.

## Acknowledgments

D.J.E. was supported by the National Institutes of Health Grant T32-GM135131.

