## Supplemental Information for "SARS-CoV-2 3CLPro Dihedral Angles Reveal Allosteric Signaling"

### Supplementary Material

#### Statistical Measures Of Dihedral Relatedness:

- Covariance: The covariance between variables $X$ and $Y$, where there are $N$ samples of each variable, is $Cov(X,Y) = \frac{\sum_{i=1}^{N}(X_i-\bar{X})(Y_i-\bar{Y})}{N-1}$. For circular/angular data, this must be modified to account for the fact that values close to $0$ are also close to $2\pi$. This was done using the equation $Cov(X,Y) = \frac{\sum_{i=1}^{N}\sin(X_i-\bar{X})\sin(Y_i-\bar{Y})}{N-1}$ where $\bar{X}$ and $\bar{Y}$ are circular means.
- Circular correlation: This is the circular version of the standard Pearson correlation coefficient. The standard Pearson $\rho_{XY}$ is the covariance divided by the product of the variables' standard deviations, $\rho_{XY} = \frac{Cov(X,Y)}{\sigma_X\sigma_Y}$. The circular correlation also uses this definition. But it uses the circular covariance and standard deviations. Here $\sigma_X$ is the square root of the variance, which is position $X$ of the diagonal of the covariance matrix.
- Inverse covariance: This is the matrix inverse of the covariance matrix.
- Mutual information: The mutual information for variables X and Y, where each can take discrete values, is $I(X;Y) = \sum_{x\in X}\sum_{y\in Y} P(x,y)\log\frac{P(x,y)}{P(x)P(y)}$ where $P$ represents probability.

Definitions for covariance, circular correlation, and inverse covariance are based on https://doi.org/10.1002/prot.26421.

| D.E. Shaw Trajectory | | |
|---|---|---|
| Dataset 1 | Dataset 2 | Pearson Between the Datasets' Dihedral Mutual Informations |
| First Third | Middle Third | 0.79 |
| First Third | Last Third | 0.56 |
| Middle Third | Last Third | 0.57 |
| First Third | Whole Trajectory | 0.79 |
| Middle Third | Whole Trajectory | 0.77 |
| Last Third | Whole Trajectory | 0.79 |
| First and Middle Thirds | Whole Trajectory | 0.90 |
| Middle and Last Thirds | Whole Trajectory | 0.90 |
| First and Last Thirds | Whole Trajectory | 0.97 |

| Random Dihedral Values | | |
|---|---|---|
| Dataset 1 | Dataset 2 | Pearson Between the Datasets' Dihedral Mutual Informations |
| First Third | Middle Third | 0.02 |
| First Third | Last Third | 0.00 |
| Middle Third | Last Third | -0.01 |
| First Third | Whole Trajectory | 0.35 |
| Middle Third | Whole Trajectory | 0.31 |
| Last Third | Whole Trajectory | 0.31 |
| First and Middle Thirds | Whole Trajectory | 0.67 |
| Middle and Last Thirds | Whole Trajectory | 0.64 |
| First and Last Thirds | Whole Trajectory | 0.65 |

*Table S1: Comparing overlapping datasets can inflate measures of consistency between datasets. For this table, we calculated the mutual informations between dihedral pairs in various samples. Then we compared how much the mutual informations from different samples agree with each other. Top: Comparisons of different thirds of the D.E. Shaw dataset. For chunks with some overlap (yellow) and lots of overlap (red), the agreement between chunks tends to be high. Bottom: the same analysis for a set of randomly generated dihedrals. Because the dihedrals are random, no agreement between different samples is expected. However, comparing two overlapping samples artificially suggests that the random dihedrals exhibit consistent, observable correlations.*

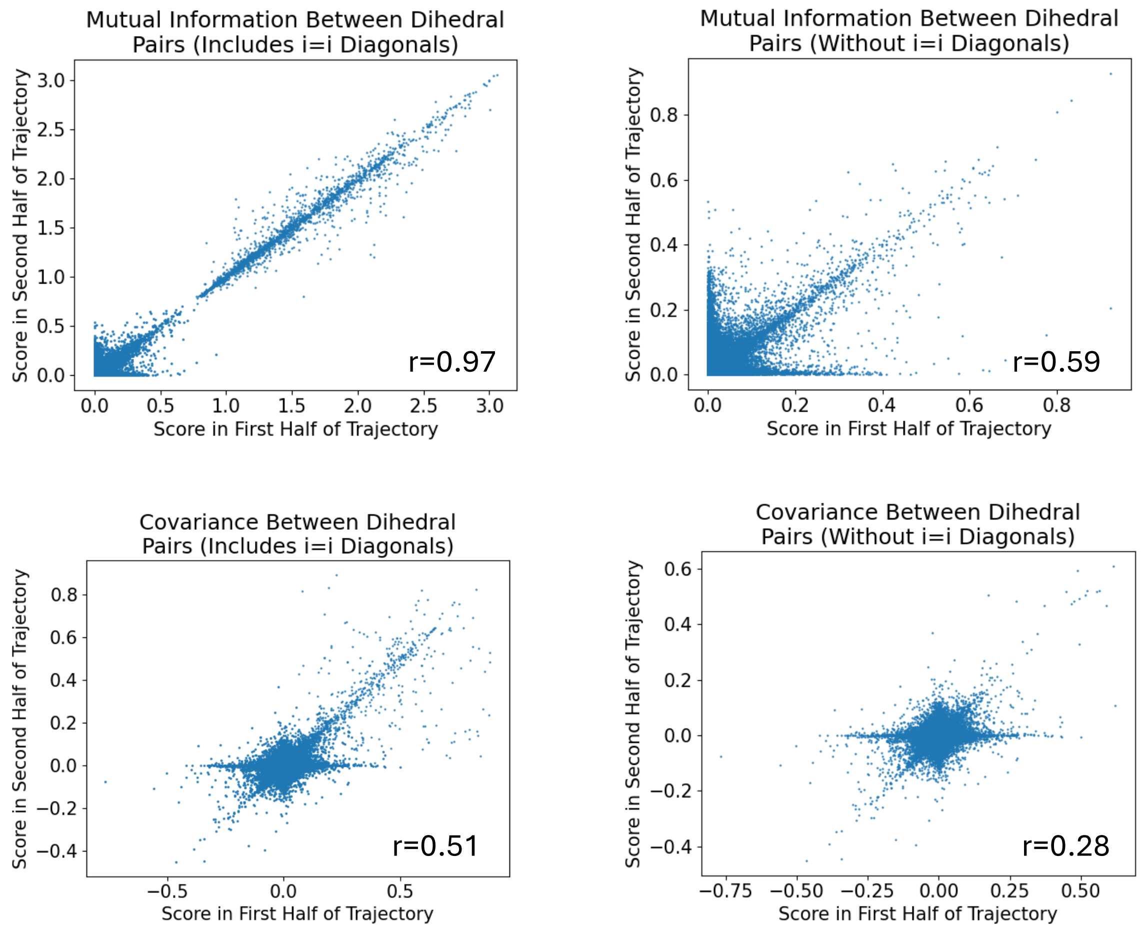


*Figure S1: Comparisons of results from the first and second halves of the D.E. Shaw trajectory. Each data point is a pair of dihedrals. The x-axes are correlation scores between dihedral pairs for the first half of the trajectory. The y-axes are correlation scores for the second half of the trajectory. The left side includes the i=i diagonal elements of the correlation matrices. This inflates the calculated agreement between scores from the two halves.*

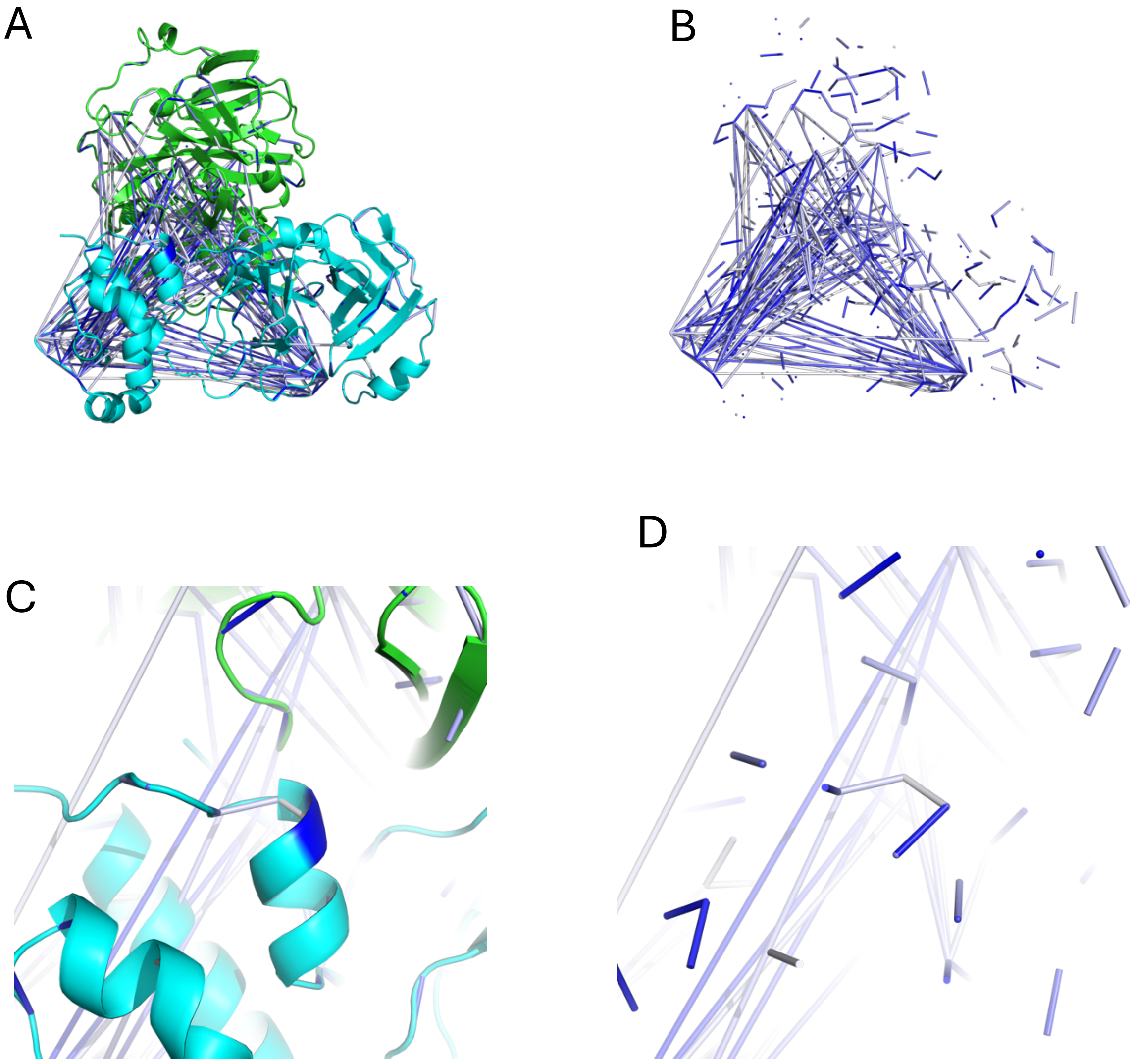


*Figure S2: The Riken simulation shows the allosteric coupling between the x1187 site and the C-terminal tail. (A) An image of the Mpro dimer. The top 0.03% of dihedral pairs, as quantified by mutual information, are shown in lines. Residue 298, which is involved in x1187 binding, is shown in dark blue. (B) The dihedral pairs from (A), but without the protein. (C) A close-up of the C-terminal tail. As in (A), residue 298 is dark blue. (D) Same as (C), but without the protein.*

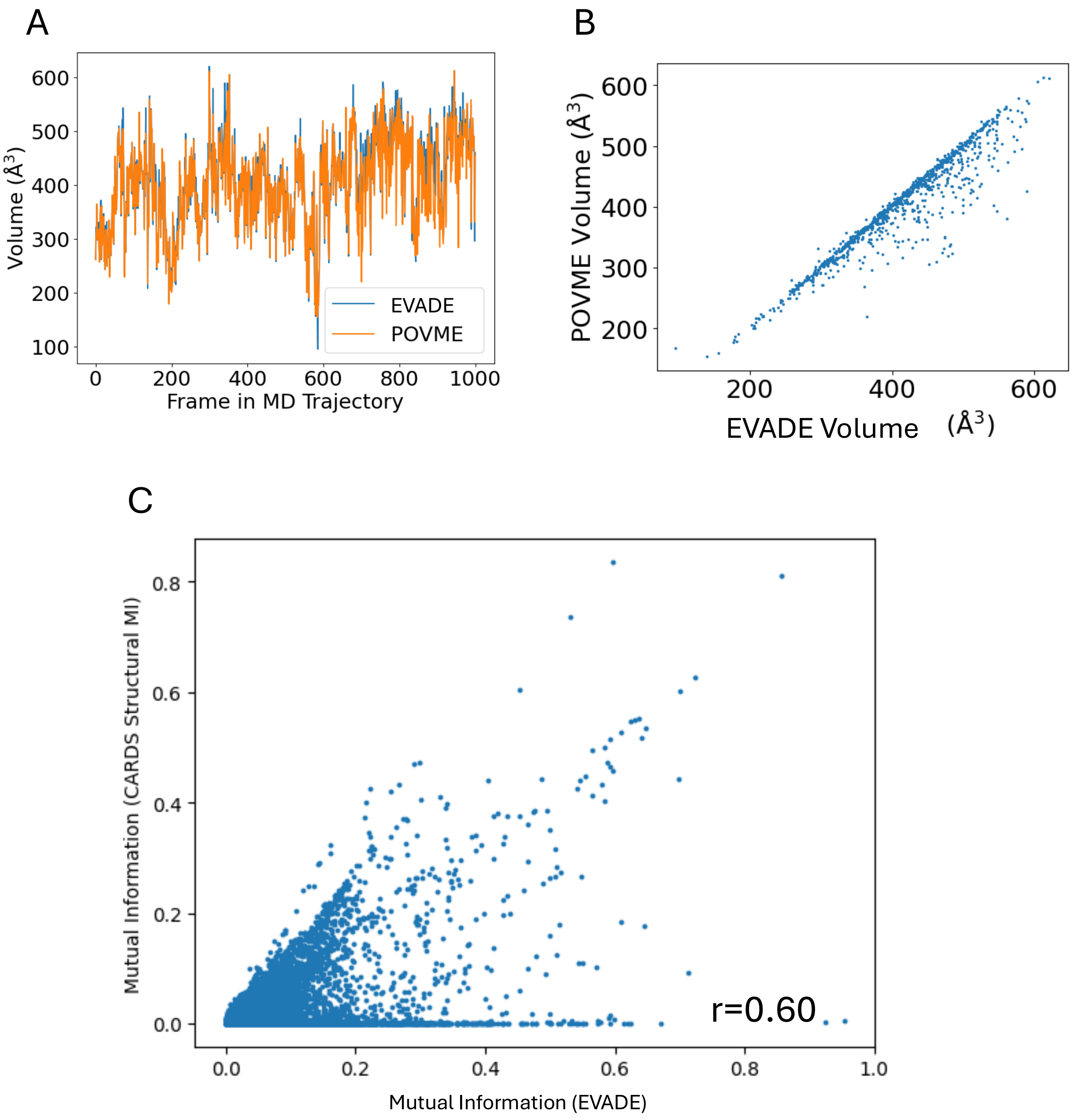


*Figure S3: Comparing EVADE with other software. (A) Pocket volumes as calculated by EVADE and POVME2. The pocket was calculated using a 9-Angstrom sphere at the center of mass of the catalytic residues (41 and 145). A solvent radius of 1.09 Angstroms was used. The volume was calculated across the first file of the D.E. Shaw dataset. (B) A scatterplot of the volumes calculated in (A). (C) Comparisons of mutual information as calculated by EVADE and CARDS. The EVADE MI was calculated with the default setting of 25 bins. The CARDS MI classifies angles into 2-3 rotameric states and does not classify changes between states unless the angle completely crosses a buffer region. We used the default setting of a 15-degree buffer region. This graph shows just the structural mutual information, i.e. not including the CARDS disorder features (since EVADE cannot perform this calculation).*
